# Normalization in mouse primary visual cortex

**DOI:** 10.1101/2023.04.18.537260

**Authors:** Zaina A. Zayyad, John H. R. Maunsell, Jason N. MacLean

## Abstract

When multiple stimuli appear together in the receptive field of a visual cortical neuron, the response is typically close to the average of that neuron’s response to each individual stimulus. The departure from a linear sum of each individual response is referred to as normalization. In mammals, normalization has been best characterized in the visual cortex of macaques and cats. Here we study visually evoked normalization in the visual cortex of awake mice using optical imaging of calcium indicators in large populations of layer 2/3 (L2/3) V1 excitatory neurons and electrophysiological recordings across layers in V1. Regardless of recording method, mouse visual cortical neurons exhibit normalization to varying degrees. The distributions of normalization strength are similar to those described in cats and macaques, albeit slightly weaker on average.

## Introduction

Nonlinearities in the responses of visual neurons, such as the saturation of the contrast response function and sublinear summation of responses when two visual stimuli are presented together, are referred to as normalization [1–4]. The responses of neurons in the visual cerebral cortex to multiple stimuli appearing simultaneously in their receptive field is generally sublinear in cats and macaques [2,5]. This normalized neuronal response to multiple stimuli typically approximates a contrast-weighted average of a neuron’s responses to those stimuli presented individually and is found across sensory modalities. Normalization also describes the large modulations of responses that occur with attentional shifts between two stimuli in a neuron’s receptive field, and may contribute to pairwise spike correlations and gamma oscillations [6–10]. The prevalence of the observation of neuronal nonlinear responses regardless of the species studied or experimental details suggests that normalization is a fundamental feature of signal processing in neural systems.

Neurons can exhibit a range of different normalization strengths, sometimes depending on the specific stimulus configuration in the receptive field [8]. Variations in the strength of normalization have been shown to correlate with attention-related modulations [6–8] and the magnitude of pairwise noise correlations [9]. This latter observation suggests that normalization arises, at least in part, from local circuit mechanisms. Consequently, to better understand normalization and its role in sensory processing, it will be important to delineate the cell types and synaptic interactions that mechanistically underlie nonlinear responses. The mouse offers many advantages for such levels of analysis, including genetic and optical access [11,12]. However, normalization is understudied in mouse sensory cortices [13,14].

To study normalization we investigated the visual response properties of V1 neurons to cross-oriented gratings using both optophysiology and electrophysiology in awake mice [2,3,15]. We find that mouse V1 exhibits visually-evoked normalization that varies in strength from neuron to neuron similar to that previously described in other species [5,7]. Moreover we find that this distribution of nonlinear responses is consistent across optophysiology and electrophysiology measurements and note that there are some differences from other species, including weaker overall strength of normalization in mice.

## Materials and Methods

### Mice

C57BL/6J (B6) mice (Jax stock #000664) and mice expressing GCaMP6s under the Thy-1 promoter (Jax stock #025776; [16,17]) were bred and maintained in accordance with the animal care and use regulations of the University of Chicago Institutional Animal Care and Use Committee. Heterozygous male and female mice (n = 12; n females = 6) aged P60-P200 were used across experiments.

### Intrinsic imaging to identify primary visual cortex

Anesthetized adult mice were surgically implanted with a head bar and a 3.0 mm or 3.5 mm diameter chronic imaging window centered over V1, as previously described [18]. Mice were anesthetized using isoflurane (1.5-2.0% in 50% O_2_) with (electrophysiology experiments) or without (optophysiology experiments) ketamine (40 mg/kg, i.p.), and xylazine (2 mg/kg i.p.).

For all experiments, a retinotopic map of the visual cortex based on intrinsic imaging was produced under anesthesia (1.25% isoflurane in 50% O_2_) two weeks after the cranial implant surgery to guide calcium imaging or electrode placement. For optophysiology experiments, intrinsic imaging was performed as in Dechery et al. [18] using drifting bar stimuli. In electrophysiology experiments, intrinsic imaging was performed as in Cone et al. [19] using stationary Gabor stimuli.

### Optophysiology

Excitatory neurons expressing GCaMP6s in Layer 2/3 of V1 were imaged using a custom-built two-photon microscope. The awake mouse was head-fixed and allowed to ambulate on a custom-made linear treadmill. Treadmill velocity was measured with a rotary encoder [18,20]. Neurons were identified by an automated algorithm [18]. A typical field of view (FOV) contained 200 to 400 identified excitatory neurons scanned at 18.6 - 28.9 Hz (mean = 23.1 Hz; SD = 3.7) [18,21]. Fluorescence data was preprocessed to calculate changes in fluorescence over baseline as previously described [18].

### Electrophysiology

Silicone multi-channel electrodes (32 sites, model 4×8-100-200-177; NeuroNexus Technologies) were used. Electrodes sampled from depths 100-1000 µm in the lateral visual field representation of V1 of anesthetized mice (1.2 - 1.5% isoflurane in 50% O_2_) approximately normal to the cortical surface. After the electrode was inserted, anesthesia was removed. Mice were awake but head-fixed and seated in a custom plastic sled during recording sessions. Signals were amplified, filtered, and sampled at threshold crossings to identify putative spikes (BlackRock Inc.) and sorted offline (Offline Sorter, Plexon Inc.) to identify small multi-unit clusters.

### Stimulus presentation

Stimuli were displayed on a calibrated video monitor positioned ∼23 cm from the mouse’s right eye during data collection. Two sets of stimuli were shown: drifting gratings in one of 8 or 12 directions to measure direction tuning and preferences, and a cross-oriented stimulus to elicit normalization.

In the optophysiology experiments, drifting square-wave grating stimuli (0.04 - 0.06 cycles/degree, 2 Hz) in eight equally spaced directions were shown in a pseudorandom order to measure the responsivity and tuning of each neuron. The drifting grating stimuli were each presented for 4 s, separated by 3 s of a luminance matched gray screen to allow time for the calcium signal to return to baseline. Each stimulus direction was shown at least 20 times.

In the electrophysiology experiments, drifting squarewave grating stimuli (0.04 cpd, 2 Hz) in 12 equally spaced directions were shown in a pseudorandom order to determine the responsivity and tuning of each neuron. The drifting grating stimuli were shown for 1 s per direction, interleaved with 1 s of a luminance matched gray screen. Each stimulus direction was shown at least 20 times.

To assess normalization, cross-oriented stimuli composed of superimposed orthogonal gratings were presented [2,15]. The cross-orientation stimuli consisted of orthogonal pairs of square-wave gratings of independently varying contrasts (0%, 6%, 12%, 25%, or 50%) superimposed to create a plaid. In six out of the eight optophysiology experiments, stimuli were counter-phased and in two the stimuli were drifting. Because responses to both types of stimuli were qualitatively similar, the data was pooled. In the electrophysiology experiments (n = 13), all stimuli were counter-phased.

### Analysis

The data was analyzed using custom-written Matlab (MathWorks) scripts. For both optophysiology and electrophysiology data, direction and orientation tuning were assessed as described previously [18,22,23]. For the optophysiology data, neurons were classified as responsive if activity evoked by a stimulus was greater than activity recorded during the preceding gray epoch for at least one grating stimulus condition. Responsiveness was established using paired sample t-tests to compare the responses of neurons to the stimulus compared with the preceding gray for each direction; p values for responsiveness were Bonferroni corrected for the number of stimulus directions (p < 0.01). Responsive units were further tested for direction or orientation tuning using repeated measures analysis of variance (p < 0.05). For the electrophysiology data, units were tested for responsiveness using an analysis of variance comparing responses to each direction with activity during gray epochs, further evaluated with Tukey’s Honest Significant Difference test (p < 0.01). Responsive units were then tested for orientation or direction tuning using a Hotelling test or t-test of the trial vectors in orientation or direction spaces (p < 0.01) [18,22]. Tuning properties of optophysiology and electrophysiology were further assessed by fitting an asymmetric Gaussian to visually evoked activity [18,22].

Activity evoked by the plaid stimulus set was calculated using the mean grating response minus the mean spontaneous activity during luminance-matched gray periods. Only neurons that were responsive to one of the grating components at 50% contrast as determined by a Kruskal-Wallis test with alpha set to 0.1 and had a minimum positive response to the normalization stimulus set were included. This minimum positive response was set to 0.1 dF/F evoked response in optophysiology experiments and 0.3 spikes/s in electrophysiology experiments.

Optophysiological and electrophysiological neuronal evoked activity to cross-oriented stimuli were analyzed using a normalization index defined as:

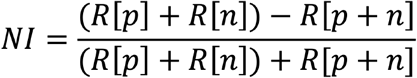

Here, R[p] is the response to the 50% contrast in the preferred plaid component, R[n] is the response to the 50% contrast in the non-preferred plaid component, and R[p+n] is the response to the 50%/50% plaid composed of both the preferred and non-preferred components.

## Results

### Optophysiological and electrophysiological recordings of mouse V1 neurons

Calcium signals were recorded from a total of 2,120 excitatory neurons from layer 2/3 in eight imaging sessions from eight sites in five awake and head-fixed mice (Fig. 1). Full contrast drifting grating stimuli were presented moving in eight directions, pseudo-randomly ordered, while imaging. Of the imaged neurons, 1,689 exhibited elevated activity to one or more gratings as compared to the response to the gray screen (Fig. 1b). In a different set of experiments and a different set of mice, multi-channel silicone probes with 32 electrode sites spanning 700 µm of V1 were inserted approximately orthogonal to the cortical surface to record single and multi-unit activity from awake, head-fixed mice. Units were recorded from 13 penetrations from seven mice. Stable recordings were obtained from 114 single units and small multi-unit clusters that had visual stimulus evoked firing rates of at least 0.3 spikes/s. Of those, 72 units were more responsive to visual stimuli than to luminance matched gray.

**Figure 1.**
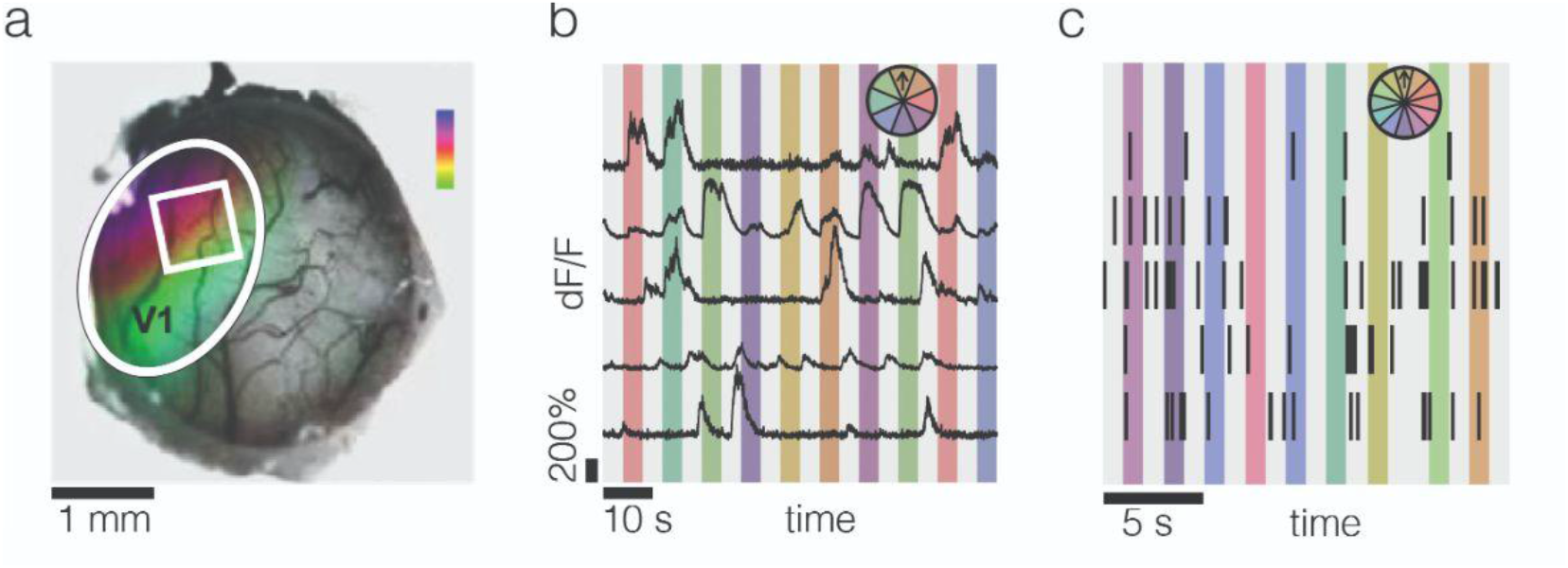
*In vivo* optophysiological and electrophysiological recordings of visual responses in mouse visual cortex. **a**, Intrinsic imaging reveals retinotopic responses in the visual cortex. The white oval represents V1; the white square represents an example two-photon imaging field of view (FOV); the color map represents azimuth from approximately 20° (violet) to 160° (green). **b**, Representative calcium fluorescence traces from five example neurons. Different colors represent 4 s presentations of eight different direction stimuli. **c**, Example spikes represented as tics recorded using multichannel silicon probes. Different colors represent 1 s presentations of twelve different direction stimuli, which were presented in pseudo-random order.

Of the responsive neurons recorded using optophysiology, most neurons were significantly tuned to orientation or direction (Fig. 2a-d). Calcium imaging signal strength can impact the calculation of normalization indices for cells with weaker stimulus responses (Supp. Fig. 2). To manage the impact of low amplitude fluorescence changes and its systematic effect on the normalization index metric, we only calculated a normalization index for cells with a minimum of a 10% increase in activity during plaid/preferred component presentation compared to the gray response. Using this criterion, a normalization index was computed for 341 imaged neurons out of the 1,211 tuned neurons. Of the responsive units in the electrophysiology experiments, 63 were direction or orientation tuned (Fig. 2e-h). Because normalization has previously been studied in relation to feature strength and selection, only tuned neurons were included in further analyses [24].

**Figure 2.**
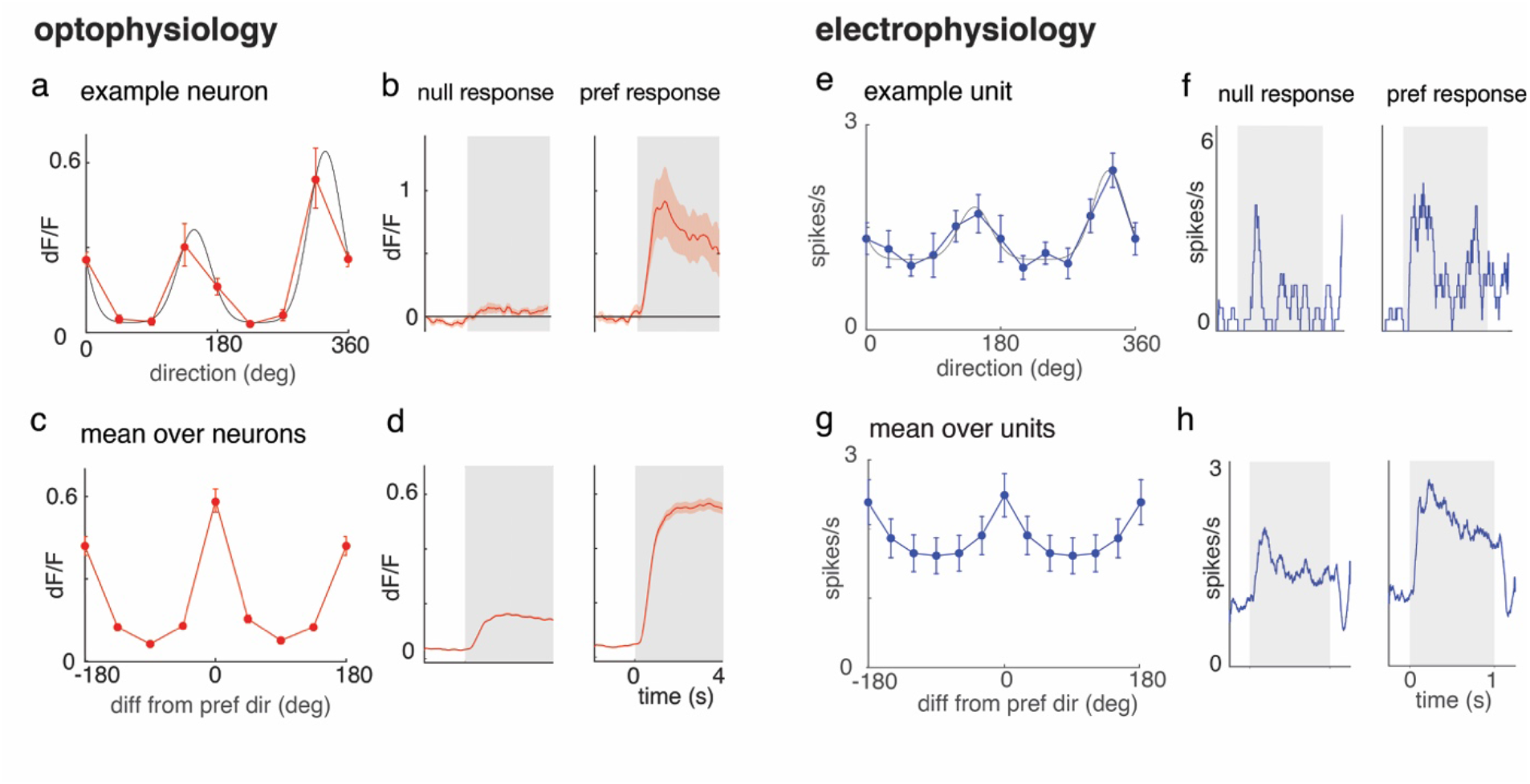
Optophysiology and electrophysiology in mouse V1 reveal direction and orientation tuned activity. **a**, An example tuning curve from a direction-selective neuron recorded optically in mouse V1. The red line is the mean dF/F of responses to drifting grating stimuli and the black line represents the tuning curve fit (r^2^ = 0.99). **b**, Mean response of the neuron shown in Fig. 2a to the drifting grating stimulus in its least preferred (null) and most preferred stimuli (pref). The gray region marks the period when the stimulus was present. **c-d**, Same as Fig. 1a and Fig. 1b, except averaged across tuned neurons (n = 341). **e**, An example tuning curve from a direction-selective unit recorded with a microelectrode in mouse V1. The blue line represents the mean response to drifting grating stimuli and the black line represents the tuning curve fit (r^2^ = 0.95). **f**, Mean response of the neuron shown in Fig. 2e to the drifting grating stimulus in its least preferred (null) and most preferred stimuli (pref). The gray region represents the stimulus-on period. **g-h**, Same as Fig. 2e and Fig. 2f, except averaged across tuned electrophysiologically recorded units (n = 63). All error bars and bands represent ±1 SEM.

### Cross-oriented stimuli elicit normalizing responses

We presented a set of counterphased cross-oriented grating stimuli [2,3,15] that included 25 combinations of orthogonal gratings at one of 5 contrasts ranging from 0% to 50% while recording from V1. To test whether these stimuli evoked normalizing responses, we used a normalization index ([7]; see methods). The index was calculated using the response to the cross-oriented (plaid) stimulus minus the linear sum of responses to the component stimuli (gratings), divided by the sum of these two quantities. For this index, a value of 0.00 corresponds to a plaid response equal to linear summation and a value of 0.33 represents a plaid response that is the average of two grating responses.

When component stimuli were superimposed to create a cross-oriented stimulus neuronal responses were normalizing (Fig 3). We found a range of normalization index values. The median normalization index as measured by optophysiology was 0.15 (interquartile range (IQR) = 0.03 - 0.25) among tuned excitatory neurons in layer 2/3 of the mouse visual cortex (Fig. 3). The bootstrapped 95% confidence interval (CI) for the median was 0.13 - 0.17.

**Figure 3.**
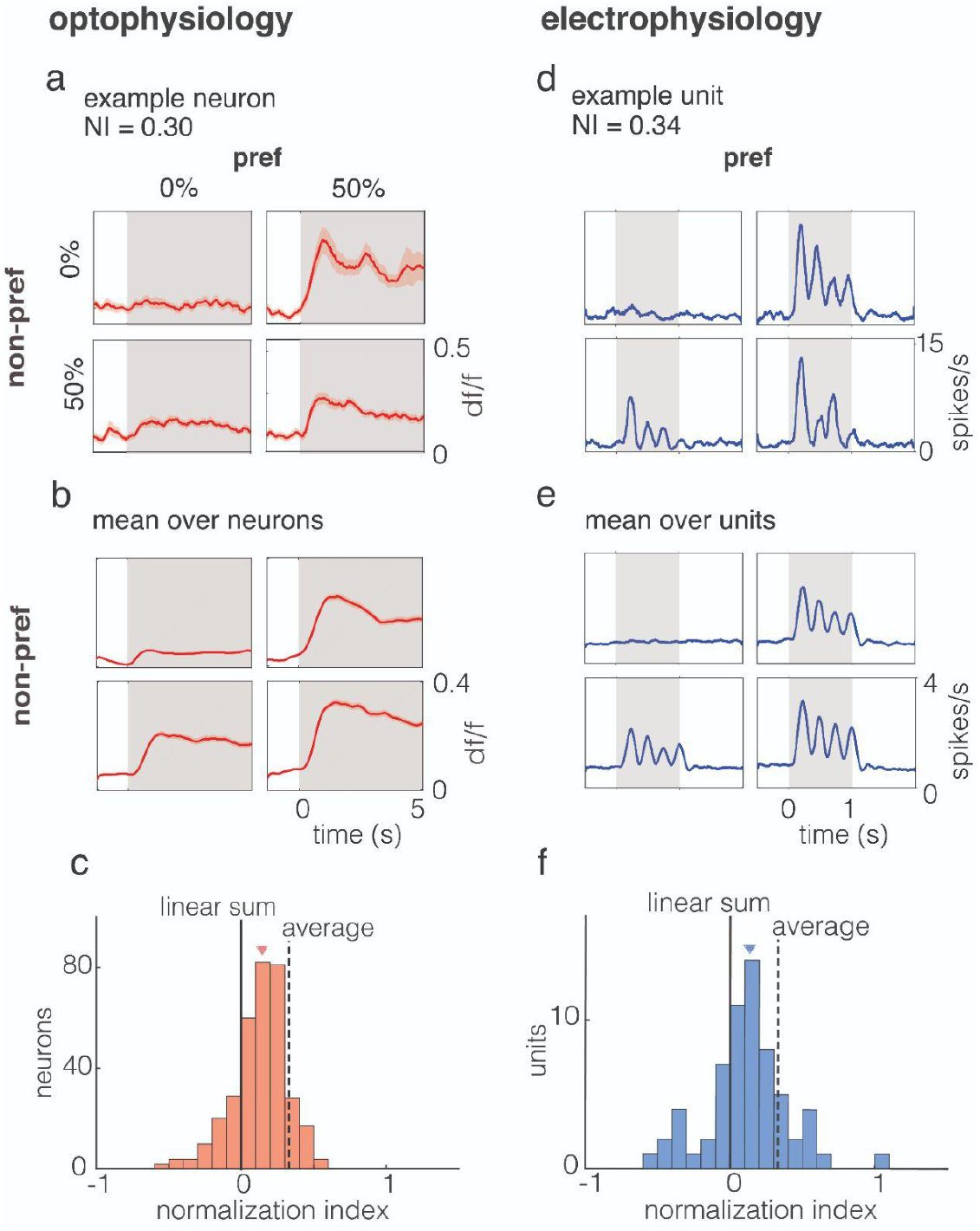
Visually-evoked normalizing responses in mouse V1 measured using optophysiology and electrophysiology. **a**, Responses of an example neuron to plaid stimuli recorded with optophysiology. Four of the 25 contrast combinations are shown. The normalization index was calculated from the evoked activity. **b**, Mean responses to plaid stimuli across neurons (n = 341). All mean response SEMs were less than or equal to 0.01. **c**, Population histogram of normalization index values (median = 0.15, IQR = 0.03 - 0.25; bootstrapped median 95% CI = 0.13 - 0.17). The median is marked with a triangle. Dashed line, expected average of normalization index (0.33) for simple averaging. **d**, Visually-evoked responses to plaid stimuli in mouse V1 recorded using electrophysiology. **e**, Mean responses to plaid stimuli across neurons (n = 63). Data points and error bars correspond to mean rate ±1 SEM. **f**, Population histogram of normalization index values (median = 0.13, IQR = −0.01 - 0.26; bootstrapped median 95% CI = 0.06 - 0.19). The median is marked with a triangle.

Mice were allowed to freely ambulate in the optophysiology experiments, and did so ∼5% of total stimulus presentation time. We found no difference in the medians or overall distribution of normalization indices regardless of inclusion or exclusion of running epochs (Supp. Fig. 1; with running: median = 0.15, IQR = 0.03 - 0.25; bootstrapped median 95% CI = 0.13 - 0.17; without running: 0.13, IQR = −0.02 - 0.25, bootstrapped median 95% CI = 0.10 - 0.15; Mann-Whitney U test, p = 0.09; two-sample Kolmogorov-Smirnov test, p = 0.16). Because running trials only composed ∼5% of total trials, this data was not analyzed separately.

When evaluating V1 neuron responses using multi-channel silicon probes, recorded units showed normalizing responses when the stimulus components were superimposed to form plaids. The median normalization index was 0.13 (IQR= −0.01 - 0.26), indistinguishable from the value measured using optophysiology. The bootstrapped 95% CI for the median was 0.06 - 0.19.

### Normalization varies with orientation selectivity

Normalization correlates with sharpness of tuning. The optophysiology data show that there is a weak but significant correlation between the width of orientation tuning and normalization (r = 0.23, p < 10^−5^). The 1-circular variance measure of the width of orientation is bounded by 0 and 1. At the extreme of 0, in which selectivity is weakest, the expected normalization index value would be approximately 0.05. At the extreme of 1, in which selectivity is greatest, the expected normalization index value would be approximately 0.23. Due to the limitations of spike sorting multi-unit clusters, the electrophysiology data could not be analyzed for circular variance.

**Figure 4.**
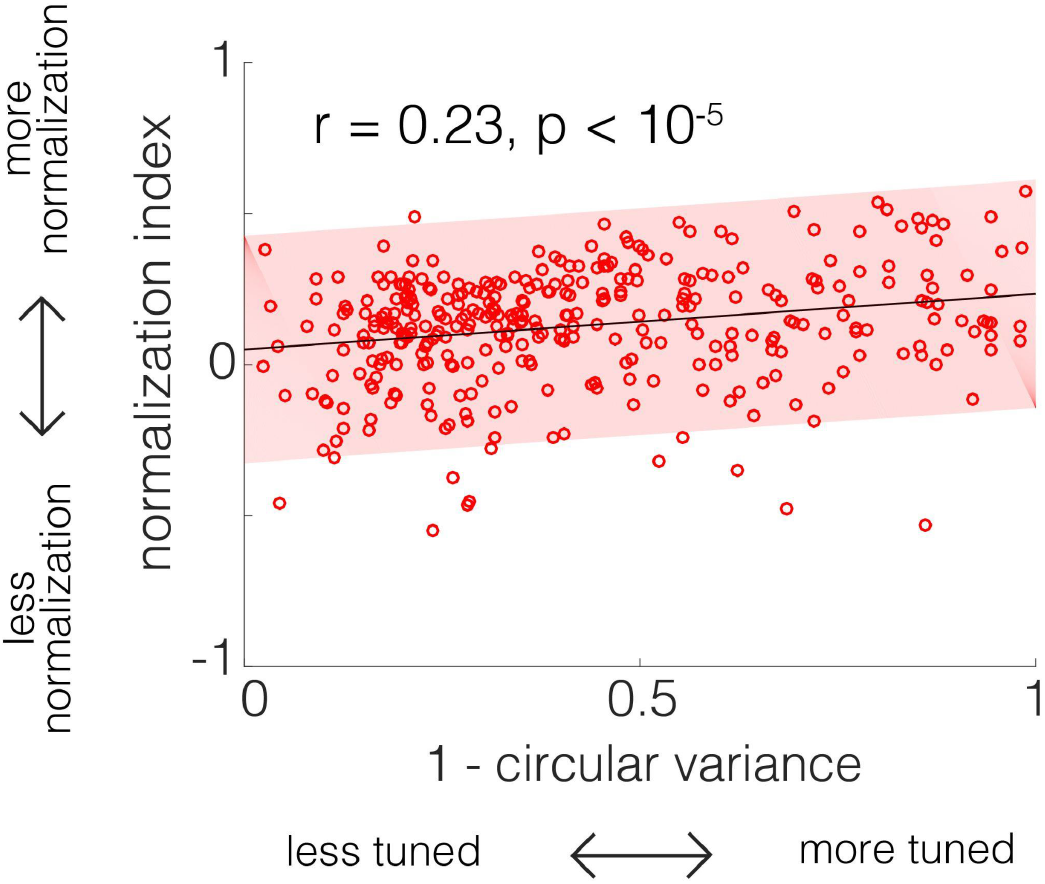
Normalization strength increases with orientation tuning strength. Optophysiological recordings demonstrate that normalization strength correlates with orientation selectivity (r = 0.23, p < 10^−5^). Linear fits and 95% prediction intervals are shown for neurons measured using optophysiology.

## Discussion

Our findings reveal visually-evoked normalizing responses to cross-oriented plaid stimuli in awake mouse V1 as measured using optophysiology and electrophysiology. The findings from the optophysiology and electrophysiology experiments were highly consistent. This suggests that the advantages of both techniques can be deployed to dissect the circuitry of normalization and that comparable measurements can be expected across these recording modalities. The range of normalization strengths is consistent with tuned normalization, in which normalization is weighted differently for different stimulus feature values (orientations) [7,9]. The overall normalization strength in mice reported here largely agrees with a recent report on the strength and range of responses to cross-oriented stimuli in the mouse primary visual cortex [13].

A recent study of macaque V1 suggests that normalization strength can be modulated by stimulus alignment with neurons’ preferred stimuli [25]. While direct comparison between these studies cannot be made due to stimulus set differences, subsets of our data that roughly correspond with that study’s analysis suggest that mouse V1 neurons exhibit weaker normalization than macaque V1 neurons. Additionally, the range of normalizing responses in mouse V1 is somewhat weaker than that reported in macaque area MT. Median normalization index values for electrophysiological measurements are ∼0.2 in macaque MT [7], compared with 0.13 for electrophysiological measures in mouse V1.

Mouse primary visual cortex neurons are overall less selective compared with macaque V1 neurons, potentially contributing to the overall weaker normalization observed in mouse primary visual cortex. Among the most selective mouse V1 neurons described here, the expected normalization index value would be approximately 0.23, similar to median normalization index values in macaque MT neurons [7]. Previous reports of tuned normalization, in which normalization within neurons was shown to vary more strongly by spatial rather than feature properties [8], are consistent with the hypothesis that less selectivity gives rise to weaker normalization. These differences in orientation selectivity and normalization could reflect differences in the cortical computational needs and strategies of different species or areas.

With the myriad tools available in the mouse, this model will facilitate future efforts to establish the cell type(s) and network structure(s) that mechanistically produce normalizing responses.

## Acknowledgements

We thank Supriya Ghosh, Elizabeth de Laittre, Yuqing Zhu, and Chery Cherian for helpful comments on drafts of this manuscript. This work was supported by NIH F30 EY030733 (ZAZ), a Naomi Ragins Goldsmith Fellowship (ZAZ), NIH R01 EY022338 (JNM), NIH U19 NS107464 (JHRM), and NIH T32 GM007281.

