## Supplemental Figures for "Normalization in mouse primary visual cortex"

**Supplemental Information**

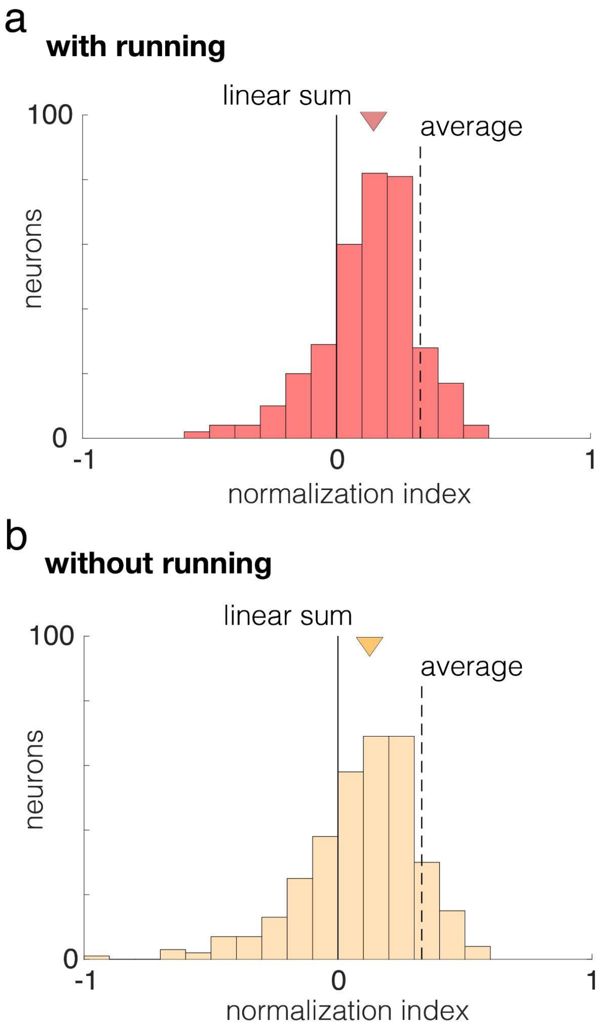


**Supplemental Figure 1. Running did not appreciably affect the median or overall distribution of normalization index**

**a,** Normalization index of neurons recorded during optophysiology experiments (n = 341). The triangle represents the median (median = 0.15, IQR = 0.03 - 0.25; bootstrapped median 95% CI = 0.13 - 0.17). **b,** Normalization index measured in the same neurons as in Supp. Fig. 1a, after excluding trials in which mice ran. The triangle represents the median (0.13, IQR = -0.02 - 0.25, bootstrapped median 95% CI = 0.10 - 0.15). No difference in the medians or overall distribution of normalization indices was statistically detectable (Mann-Whitney U test, n.s.; two-sample Kolmogorov-Smirnov test, n.s.). A normalization index of 0.00 represents the expected normalization index for linear summation and the dashed line at 0.33 represents the expected normalization index for simple averaging.

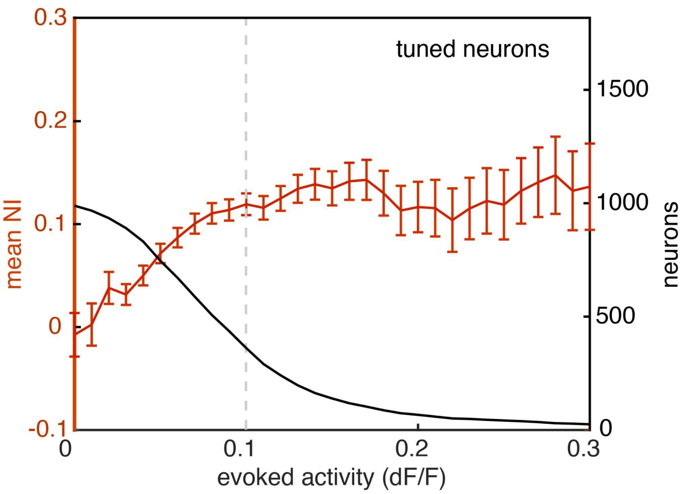


**Supplemental Figure 2. Visually-evoked normalization varies systematically with calcium signal strength**

The red line shows the normalization index as a function of minimum evoked response to the preferred and plaid stimulus over the gray condition in tuned neurons. As the evoked activity of tuned neurons approaches zero, the normalization index decreases due to erosion in the signal to noise ratio. The black curve shows the number neurons that are included at each threshold level. The number of neurons included at each threshold level decreases as the minimum evoked activity increases. The dashed gray line represents the threshold selected. Error bars indicate mean ± 1 SEM.
